# Multiple forms of working memory emerge from synapse-astrocyte interactions

**DOI:** 10.1101/2021.03.25.436819

**Authors:** Maurizio De Pittà, Nicolas Brunel

## Abstract

Competing accounts propose that working memory (WM) is subserved either by persistent activity in single neurons, or by time-varying activity across a neural population, or by activity-silent mechanisms carried out by hidden internal states of the neural population. While WM is traditionally regarded to originate exclusively from neuronal interactions, cortical networks also include astrocytes that can modulate neural activity. We propose that different mechanisms of WM can be brought forth by astrocyte-mediated modulations of synaptic transmitter release. In this account, the emergence of different mechanisms depends on the network’s spontaneous activity and the geometry of the connections between synapses and astrocytes.

---

The neural basis of working memory (WM) remains an open problem. A large body of evidence supports a role for sustained neural activity in prefrontal and other cortices, possibly supported by attractor dynamics in recurrently connected circuits (Constantinidis et al., 2018). In this view, neurons hold sensory information beyond the presentation of a sensory-relevant cue by their persistent firing activity (PA). However, competing evidence suggests that WM could also be accounted for by dynamically varying activity patterns and activity-silent representations (Kamiski et al., 2020).

There is a growing debate on whether different mechanisms of WM could coexist within the same brain region (Barbosa et al., 2020), and the underpinning cellular substrate for their coexistence remains elusive (Kamiski et al., 2020). Activity-dependent synaptic facilitation has emerged in recent years as an appealing candidate mechanism for this coexistence (Barak et al., 2014; Mongillo et al., 2008). On the one hand, it bestows cortical networks with slow time scales (from hundreds of milliseconds to minutes) that might help stabilize PA (Barak et al., 2014; Chaudhuri et al., 2016). On the other, it can also encode memory by synaptic variables whose dynamics can sustain WM in the absence of PA (Mongillo et al., 2008). Although synaptic facilitation is traditionally regarded as a purely neuronal process, we consider here the possibility that it could also involve glial signaling.

Among cortical glial cells, astrocytes are ubiquitous in the neuropil. They are prominently found in proximity of nerve terminals, sensing neural activity and being activated during synaptic transmission. Astrocytes can also modulate synaptic transmission by releasing transmitters – dubbed “gliotransmitters” for their glial origin – like glutamate (Araque et al., 2014). In particular, gliotransmission may promote synaptic release from excitatory terminals for several seconds, up to minutes, thus potentially contributing to WM processing, akin to short-term synaptic facilitation, but on longer time scales. In agreement with this hypothesis is the observation that astrocyte stimulation in mice’s primary sensory areas increases neural firing beyond stimulation (Perea et al., 2014; Poskanzer et al., 2016). This phenomenon appears in association with changes of neuronal gain by higher concentrations of extracellular glutamate, possibly due to the enhanced release of this neurotransmitter by gliotransmission (Perea et al., 2014). In this fashion, gliotransmission would mediate a positive feedback on neuronal activity that could also be relevant for WM processing (Chaudhuri et al., 2016).

To investigate this possibility, we started by analyzing a minimal neuron-glial circuit (Figure 1a) of a single integrate-and-fire neuron stimulated by *N* synapses. A fraction *f* of those synapses are shared with an astrocyte, leading to interactions in both directions (Figure 1b). Incoming action potentials (APs) trigger synaptic release, which occur stochastically, *r* = 1, with probability *u*, and 0 otherwise (Figure 1c). An integrate-and-fire formalism is also adopted to describe astrocyte activation (*ν_G_*) in terms of slow build-up and fast release of intracellular calcium mediating gliotransmitter release (Online Methods). When *ν_G_* = 1, a calcium ‘spike’ occurs, triggering glutamate release from the astrocyte with some probability (Figure 1d). Each gliotransmitter release event (GRE), in turn, transiently increases glutamate release probability at those synapses shared between the astrocyte and the neuron (Figure 1e). In agreement with experimental observations (Araque et al., 2014), this increase decays slowly, with a time scale *τ_G_* > 5 s, to the value of release probability attained by the synapses in the absence of gliotransmission (SOM).

**Figure 1.**
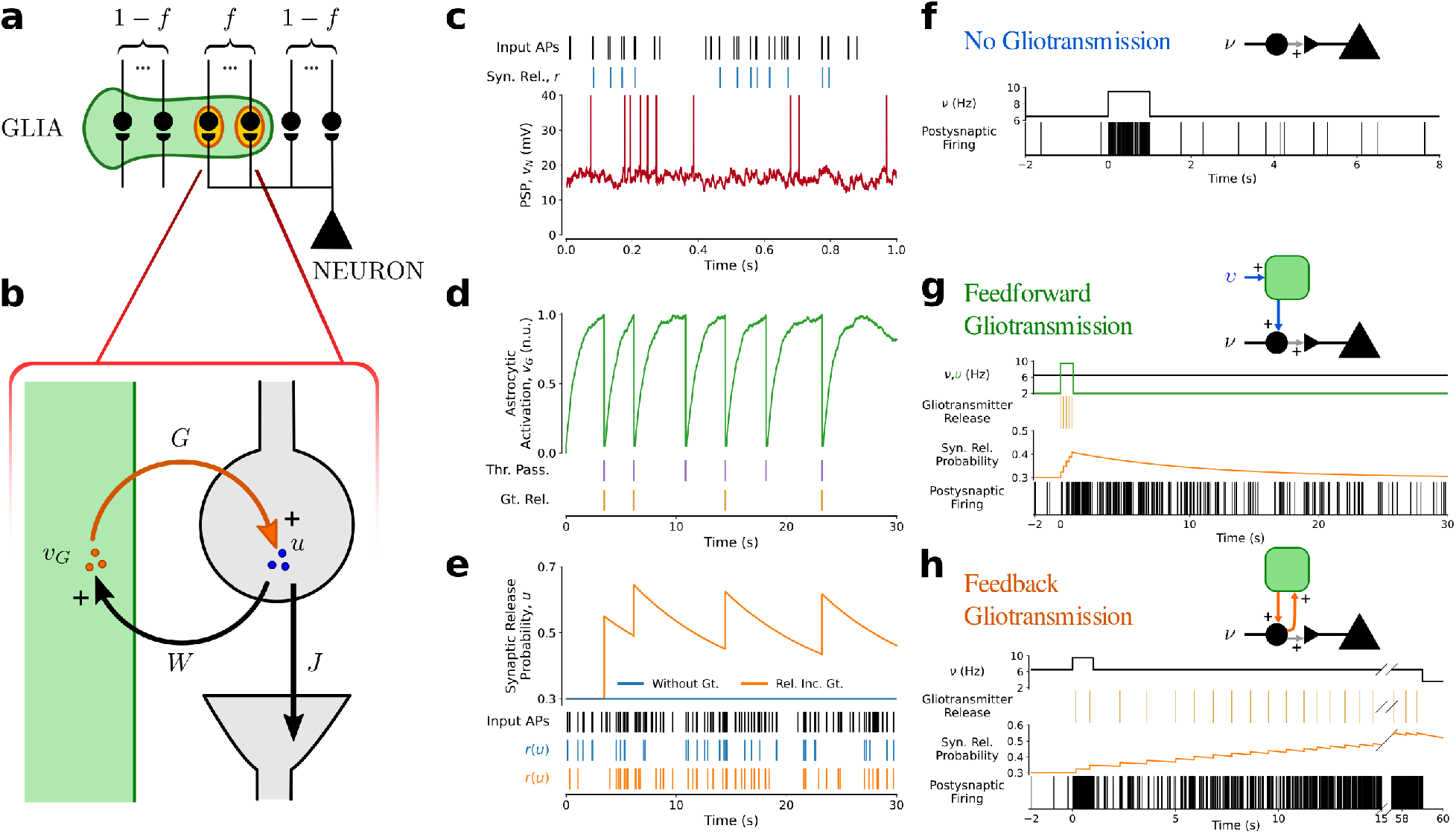
Minimal neuron-glial circuit. (**a**) Schematics of a single neuron-astrocyte domain, where a fraction *f* of the synapses are shared by the neuron with the astrocyte. (**b**) Synapse-astrocyte positive feedback loop. Neurotransmitter release *r* at a presynaptic terminal, occuring with probability *u*, can depolarize both the post-synaptic neuron (by an amount *J*), and the astrocyte (*W*). Astrocyte activation can trigger glutamate release from the astrocyte which, in turn, can lead to an increase in release probability *u* described by a parameter *G*. (**c**) Incoming action potentials at a sample synapse (*black raster*) trigger stochastic neurotransmitter release (*blue raster*). The summed synaptic input from *N* = 1000 excitatory synapses drives fluctuations in the membrane potential (*ν_N_*) of a leaky integrate-and-fire neuron (*red traces*), leading to irregular firing (*vertical red bars*). (**d**) The shared synapses also stimulate the astrocyte whose activation (*ν_G_*) is equivalently described by a leaky integrate-and-fire formalism (*green trace*). Each calcium ‘spike’ (*purple vertical lines*) triggers stochastic glutamate (Gt.) release from the astrocyte (*yellow vertical lines*). (**e**) The instantaneous neurotransmitter release probability from a synapse modulated by astrocytic glutamate (from **d**) (*orange trace*) is shown together with the release probability from the same synapse in the absence of gliotransmission (*blue line*). *Bottom rasters* exemplify how these two scenarios results in different transmission of a train of APs (*black raster*). (**f**) In the absence of gliotransmission, the circuit is memoriless, and neuronal firing (*raster*) quickly returns to baseline right after the presentation of a square-pulse input current ((*top panel*, for 0 ≤ *t* < 1 s). (**g**) Independent stimulation of the astrocyte (*green square pulse*), to trigger gliotransmitter release therefrom (*yellow raster*) results in the transient increase of synaptic release probability (*orange trace*) in association with a long-lasting transient increase of postsynaptic firing (*black raster*). (**h**) Postsynaptic firing can ramp up, and eventually turn persistent, when gliotransmission is stimulated by the same synapses that it modulates (*yellow raster*). When this occurs, synaptic release is bistable, leading to two stable post-synaptic firing rates, one low, the other high (*black raster*, for *t* < 0 and *t* > 15 s).

In the absence of the astrocyte, this minimal circuit is memoryless - the neuronal firing rate only depends on the current inputs, and while an increase in input can increase the neuron’s firing rate, this rapidly goes back to baseline after the original input is restored (Figure 1f). In the presence of gliotransmission, the neuronal response to an input can change dramatically, with two possible scenarios. In Figure 1g, gliotransmission is triggered by stimulating the (1 – *f*)*N* synapses that are not shared with the neuron, while the stimulation rate of the neuron remains constant. In this fashion, gliotransmission occurs only during the cue presentation (*yellow marks* coinciding with the *green square pulse* in the top panels), yet it promotes a slow-decaying increase of neurotransmitter release at synapses shared between the neuron and the astrocyte (*orange trace*). This increases the net synaptic drive to the neuron, resulting in a transient increase of its firing activity that decays over time scales of the order of *τ_G_* (*bottom raster*).

A ramping activity can be obtained when gliotransmission is triggered by synapses that also stimulate the neuron (Figure 1h). In this case, the positive feedback of gliotransmission on synaptic stimulation can promote astrocytic glutamate release beyond the cue’s presentation and, in turn, keep higher levels of synaptic release in a self-sustained manner. For sufficiently strong astrocytic activation, PA may also emerge following the cue, even though presynaptic stimulation recovers to pre-cue rates, thanks to the emergence of bistability (SOM). This bistability is due to the positive feedback loop involving gliotransmission: the synaptic drive to the neuron for a given input rate can be high or low depending on whether release-increasing gliotransmission is occurring or not (Figure S2). In particular, when an input pushes synaptic release from the low activity stable attractor to the high one, the neuron firing ramps up after the cue presentation, similar to experimental observations in delay periods of delayed response tasks (Barbosa et al., 2020; Stokes, 2015; Inagaki et al., 2019). Neuronal firing ultimately reaches a constant steady-state value until a sufficiently long decrease in external input resets the neuronal firing rate to baseline.

Bistability of synaptic release and neuronal firing by feedback gliotransmission in the minimal circuit occurs robustly for a range of neuronal and astrocytic parameters, accounting for various possible neuron-glial ensembles in the brain (Figure S3). The next question we ask is whether bistability could also emerge in large cortical neuron-glial networks. To answer this question, we considered a network of 4000 excitatory (E) and 1000 inhibitory (I) randomly connected neurons (Brunel, 2000), that interact with 4000 astrocytes (G). Motivated by experimental data, both excitatory and inhibitory synapses stimulate astrocytes, but only excitatory ones are modulated by gliotransmission (Araque et al., 2014) (SOM). Analysis of the asynchronous state of the EI+G network constructed by randomly assigning recurrent synapses to the astrocytes with equal probability confirms the possibility of co-existing UP and DOWN states for fixed global unspecific inputs (Figure S4). In particular, bistability emerges in the inhibition-dominated regime by feedback glio-transmission, providing a mechanism for WM in a network that would be otherwise memoryless without astrocytes. Moreover, in such a regime, the increase of excitatory drive mediated by glio-transmission competes with the strong inhibition, leading to firing rates of the UP state that can differ by just a few Hz from the DOWN state (Figure S3). In this way, a short high-frequency step increase of afferent stimulation can push the network from the DOWN to the UP state, triggering PA at a rate that is close to background rate (Figure 2a). At the same time, ongoing gliotransmission changes the internal state of excitatory synapses, providing a latent mechanism for WM by facilitation of synaptic release probability (*u*, *orange trace*). This also holds for more silent versions of WM achieved by lower spontaneous activity and allows reactivating the cue’s memory by weaker nonspecific inputs presented to the network, as long as gliotransmission-mediated synaptic facilitation is high enough (Figure S6).

**Figure 2.**
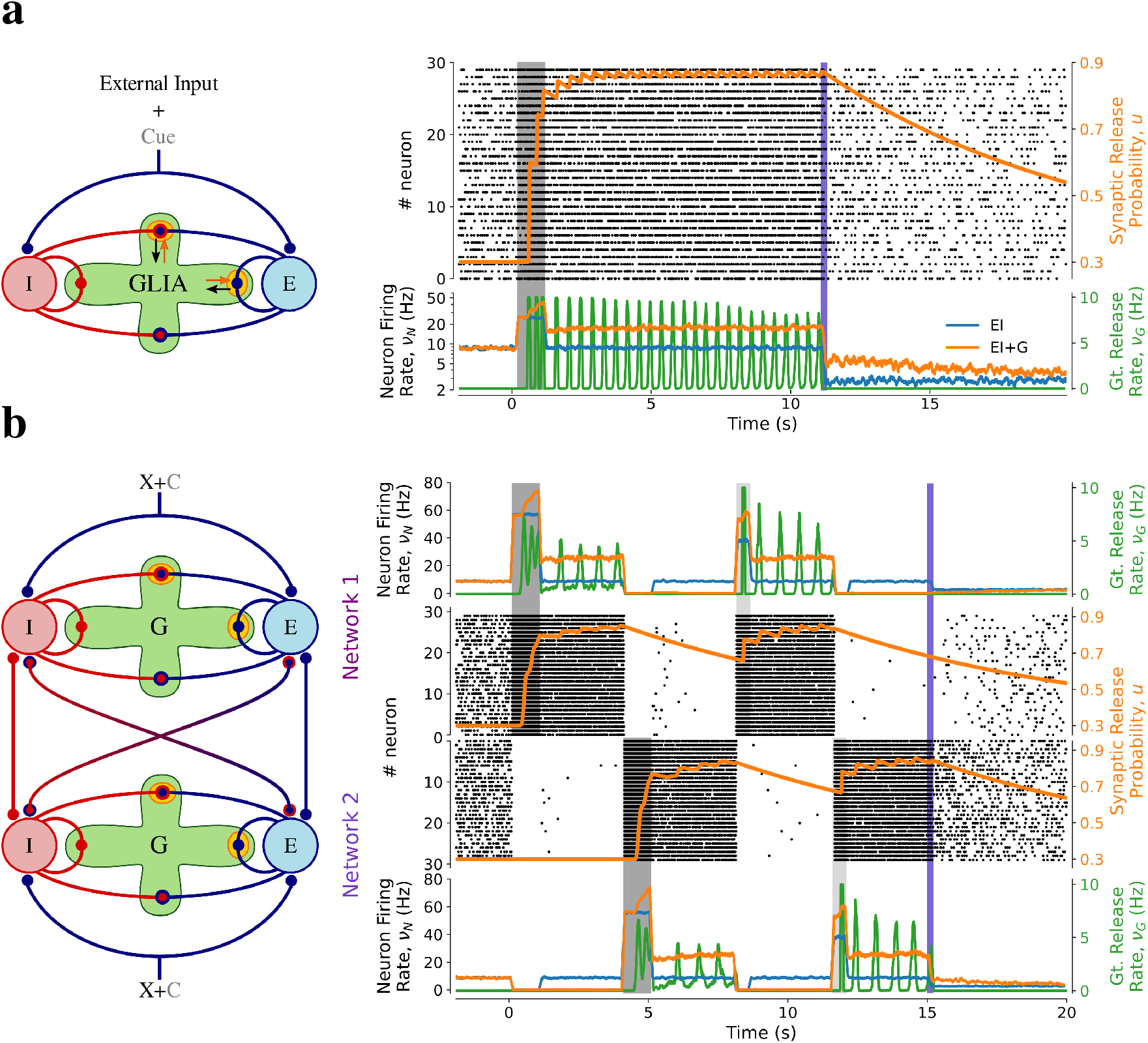
Mechanisms of working memory in neuron-glial networks. (**a**) Random neuron-glia network. The network is stimulated by a brief square pulse (cue ‘C’ with rate of ~25 Hz) for 0 ≤ *t* < 1 s (*dark shading*). Without gliotransmission, neuronal firing quickly recovers to baseline after cue presentation (*blue trace*). In the presence of gliotransmission, the cue promotes a persistent increases of synaptic excitation (*u*, *top orange trace*), resulting in PA. *Black dots*: spike rasters of a representative subset of 30 excitatory neurons. *Purple bar* marks a reduction of afferent excitation, leading to a reduction in neuronal activity. (**b**) Multistability in networks composed of two distinct neuron-glia subnetworks, encoding two distinct memories. Neuronal connectivity is unstructured, but each astrocyte population modulates only recurrent synapses within its own sub-network. A random subset of 50% of neurons in subnetwork 1 is stimulated by a high-frequency (~60 Hz) cue, promoting PA in subnetwork 1, while suppressing most of the subnetwork 2’s activity thanks to recurrent inhibition. At *t* = 4 s, the cue is fed to the subnetwork 2, making it persistently active, while suppressing activity in subnetwork 1. Individual memories can be reactivated by weaker stimulations (30 Hz) of single subnetworks, occurring within a temporal window of about *τ_G_* = 10 s.

The topology of synapse-astrocyte interactions is an additional important factor in shaping network dynamics. Different synaptic ensembles modulated by distinct astrocytes could promote discrete domains of excitation and inhibition (Halassa et al., 2007), resulting in heterogeneous active neuronal populations. Consider the scenario in Figure 2b where the astrocytes in the EI+G network belong to two disjoint populations that split the neurons into distinct subnetworks whose recurrent connections are modulated by distinct astrocytes. Without astrocytes, the selective stimulation of a subnetwork transiently promotes the firing of neurons in that subnetwork that also leads to a transient surge of feedback inhibition to the rest of the network, suppressing the firing of unstimulated neurons (*blue traces* in the *dark gray shades*), as in standard attractor neural networks with strong recurrent inhibition (e.g. (Brunel and Wang, 2001)). Moreover, after cue presentation, no PA is observed as the network is dominated by inhibition. With the addition of astrocytes, the stimulated subnetwork can instead turn persistently active by the selective stimulation of the associated astrocyte population. In this way, the ensuing increase of recurrent excitation promoted by gliotrasmission remains spatially confined within the subnetwork (*top u orange trace*). The whole network can also be switched to another ‘memory state,’ by a sufficiently strong stimulation of the second subnetwork, which leads to suppression of the first (*gray shade* at *t* = 4 s). Moreover, thanks to the slow time scale τ_G_ of decay of the modulation of synaptic excitation by gliotransmission, it is possible to reactivate individual memories by ~30% weaker stimuli to individual subnetworks (*light gray shades*). In this fashion, different domains of gliotransmission from distinct astrocyte populations, promote clustered network activity. Each cluster encodes a separate WM item, and emerges as a neuronal population that is kept persistently active by spatially-segregated astrocyte activation, and the associated gliotransmission modulating synaptic release.

We propose that modulations of synaptic transmission by gliotransmission could participate in sustaining different working memory-related patterns of activity observed experimentally, in which neuronal firing can ramp up and turn persistent after a cue. Alternatively, neural activity can also ramp down to pre-stimulus levels past the cue, in which case the memory item is encoded by changes of synaptic weights, that are sustained by gliotransmission. This mechanism can potentially lead to very low levels of delay activity, which could be consistent with ‘silent’ working memory phenomena reported experimentally (Schneegans et al., 2017). It will be important to characterize further experimentally the anatomy of the connections between cortical synapses and astrocytes (Santello et al., 2019), given our observation that multistability could emerge from neuron-glia networks in which astrocytes define specific neuronal clusters. In light of the emerging heterogeneous astrocyte arrangement across the brain (Lanjakornsiripan et al., 2018), future efforts should aim to characterize the variety of possible neuron-glial networks’ attractors and their entire dynamical repertoire.

## Acknowledgements

We thank Paolo Bonifazi for comments on the manuscript, and Marcel Stimberg for technical support on the code to simulate the spiking models. This work was funded by a Junior Leader Postdoctoral Fellowship to MDP by ‘la Caixa’ Banking Foundation (LCF/BQ/LI18/11630006).

## Online Methods

### Computational models

#### Single neuron-astrocyte domain

We consider a minimal neuron-glial circuit of a neuron (N) and an astrocyte (G). The neuron has *N* = 1000 identical synapses in total feeding into the circuit, that are independently stimulated by trains of Poisson-distributed action potentials (APs), modelled by Dirac delta fnctions, i.e., 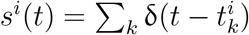 for the *i*th synapse. A fraction *f* of these synapses is “shared” by the neuron and the astrocyte. Those synapses stimulate both cells, and are modulated by gliotransmission from the astrocyte. Glutamate release (*r*) from synaptic terminals is described by a Bernoulli process: when an AP reaches a presynaptic terminal, *r* = 1 with probability *u* for release success, and *r* = 0 otherwise. If successful, glutamate release instantaneously depolarizes the neuron by *J*, while increasing astrocytic activation by *W*. Neuronal depolarization (*ν_N_*) is described by a leaky integrate-and-fire (LIF) formalism, such that

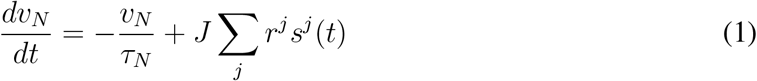

where *τ_N_* is the characteristic decay time constant for neuronal depolarization. Every time *ν_N_* reaches a threshold value 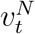, the neuron generates an AP, *ν_N_* is reset to 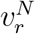, and subsequently held to this value for a refractory period 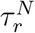.

We also consider a LIF description for astrocytic activation. This activation can be thought as being an increasing function of the astrocyte’s intracellular Ca^2+^ concentration, which critically regulates gliotransmitter release (SOM). Astrocytic Ca^2+^ activity (*ν_G_*) is driven by synaptic stimulation *i_G_*(*t*) according to

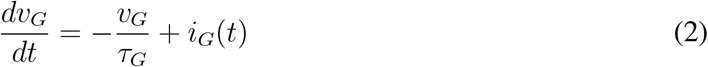

with 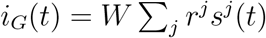, and *τ_G_* is a lumped time constant for stimulus integration by astrocytic Ca^2+^ signaling (Supplementary Text Section 1.1). Because gliotransmitter release only occurs when Ca^2+^ reaches a threshold 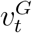, a gliotransmitter release event (GRE) is set to occur when 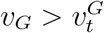, following which *ν_G_* is reset to and held at baseline 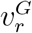 for a refractory period 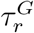 (SOM).

Because both astrocytic calcium signaling (Semyanov et al., 2020) and gliotransmitter release have a stochastic component (Potokar et al., 2013; Bowser et al., 2007), we also consider a stochastic description of GREs. That is, GREs are modelled by a Bernoulli random variable *r_G_*, which is 1 (release success) with probability *u_G_*, and 0 (release failure) otherwise. The single gliotransmitter release event in this context represents the ‘total release’ of gliotransmitter per threshold crossing, and it does not account for potentially different mechanisms of gliotransmitter release (Savtchouk et al., 2018). In this fashion, the sequence of GREs originating from the astrocyte, each occurring at instants 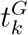, is described by 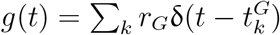.

At synapses that are shared with the astrocyte, synaptic release probability *u* is taken to be modulated by GREs from the astrocyte according to (Supplementary Text Section 1.2)

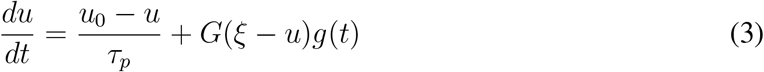

where *u_0_* stands for the (baseline) synaptic release probability in the absence of gliotransmission; *G* and > *u*_0_ control the strength of the positive feedback of gliotransmission on synaptic release, and *τ_p_* is the decay time constant of the increase of synaptic release mediated by gliotransmission.

#### Neuron-glial network

The model network of excitatory and inhibitory (EI) originally introduced by Brunel is extended to include astrocytes. The network is composed of *N_G_* = 4000 astrocytes (G), together with *N_E_ = 4000 excitatory neurons (E) and *N*_I_* = 1000 inhibitory neurons (I). All neurons (astrocytes) are described by a LIF formalism, with identical membrane time constants *τ_N_* (*τ_G_*), and firing thresholds 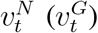. Neurons in the network are randomly connected such that each neuron receives *C_E_* = 400 connections from excitatory neurons, and *C_I_* = 100 connections from inhibitory neurons. Additionally, all neurons are supposed to be stimulated by *C_E_* excitatory afferents from the same cell population (X) outside the network. Synaptic release (*r*) at recurrent excitatory connections is probabilistic, with a probability that depends on GREs from the astrocytes, as described above. For simplicity, we consider identical postsynaptic potential amplitudes *J* > 0 at excitatory synapses, and –*g_J_J* at inhibitory synapses. The external synapses are taken to be stimulated by independent Poisson processes with rate *ν_*X*_ = ρνθ*, where 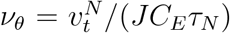 is the external frequency that leads to an average membrane potential equal to the neuronal firing threshold. In this fashion, the external input to the network is modelled by a background noisy current, with mean *μ_X_* = *C_E_Jν_X_τ_N_* and variance 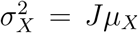. Denoting by 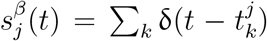 the train of incoming action potentials from neuron *j* in population *β*, the depolarization 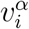 of the *i*th neuron in population *α* evolves according to:

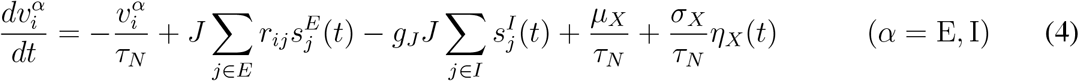

where *η_X_*(*t*) is a temporally uncorrelated normal random variable with mean 0 and variance 1.

All synapses, regardless of their type, can depolarize glial cells. This reflects the experimental observation, that both glutamatergic (excitatory), and GABAergic (inhibitory) synapses can trigger calcium-dependent gliotransmission (Araque et al., 2014; Losi et al., 2014). We assume that all synapses contribute equally to astrocytic depolarization by *W*. Moreover, given that the molecular machinery for gliotransmission is found in juxtaposition with synaptic terminals (Jourdain et al., 2007), we assume that each excitatory synapse stimulating an astrocyte, is also modulated by gliotransmission from the latter (Supplementary Text Section 2.2). Hence, the depolarization vG of the i-th astrocyte and the release probabilities of excitatory synapses are described by the set of coupled differential equations

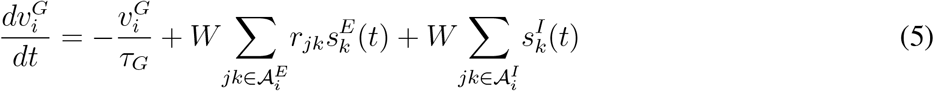

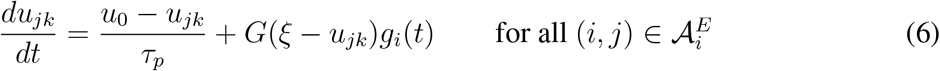

where *g_i_* is the train of GREs originating from the *k* astrocyte, and 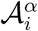 is the set of all *α* synapses impinging on astrocyte *i*. Specifically, we consider different scenarios for connections between recurrent synapses and astrocytes. In the random network of Figure 2a, each astrocyte picks a random subset *C*/*N_G_* of synapses, out *C* = (*C_E_* + *C_I_*)(*N_E_* + *N_I_*) recurrent synapses. In the network of Figure 2b instead, the EI network was partitioned into two subnetworks of equal size, in terms of cell numbers and number of synapses (i.e. *C*/2). In this case, the recurrent connections of each subnetwork were assigned to distinct populations of *N_G_*/2 astrocytes.

### Mathematical analysis

Mean-field analysis of the minimal neuron-glial circuit and the unstructured EI+G network (Figure 2a) was performed along the lines of (Amit et al., 1997). Details can be found in the Supplementary Text Section 3. Briefly, the mean-field analysis yields self-consistent equations for the average rates of gliotransmitter release (*ν_G_*) and firing rates of both E and I populations. For our parameter choice, the mean firing rate of excitatory neurons equals that of inhibitory cells, *ν_E_* = *ν_I_* = *ν_N_*. Both *ν_N_* and *ν_G_* rates depend on the first two moments of the synaptic inputs to neurons and astrocytes, that in turn can be expressed as functions of these rates, and the average synaptic release probability at excitatory synapses *U*(*ν_G_*). In this way, the mean rates are solutions of the equations

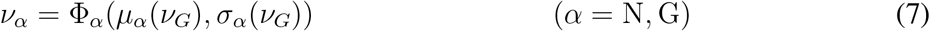

where Φ_*α*_(*μ_α_*, *σ_α_*) yields the firing rate of a cell in population *α* receiving noisy inputs of mean *μ_α_* and variance *σ_α_*.

### Numerical methods

Simulations and mean field analysis used custom code implemented in C/C++, Python 3.x, and the Python-based Brian 2.x simulator (Stimberg et al., 2019). Numerical integration used either an event-based scheme, or the Euler-Maruyama method with time step *dt* = 10 μs. Bifurcation diagrams were generated by Matcont 6.x in Matlab^®^ (R2017a, MathWorks) (Dhooge et al., 2008).

Simulations were performed on an Intel^®^Xeon^®^ 12-core CPU E5-1660 @3.30 GHz Linux desktop. The Python classes to reproduce the neuron-glial circuits of this study are available from the authors upon request.

